## Supplementary materials for "Perceived and mentally rotated contents are differentially represented in cortical depth of V1"

### Supplementary Information

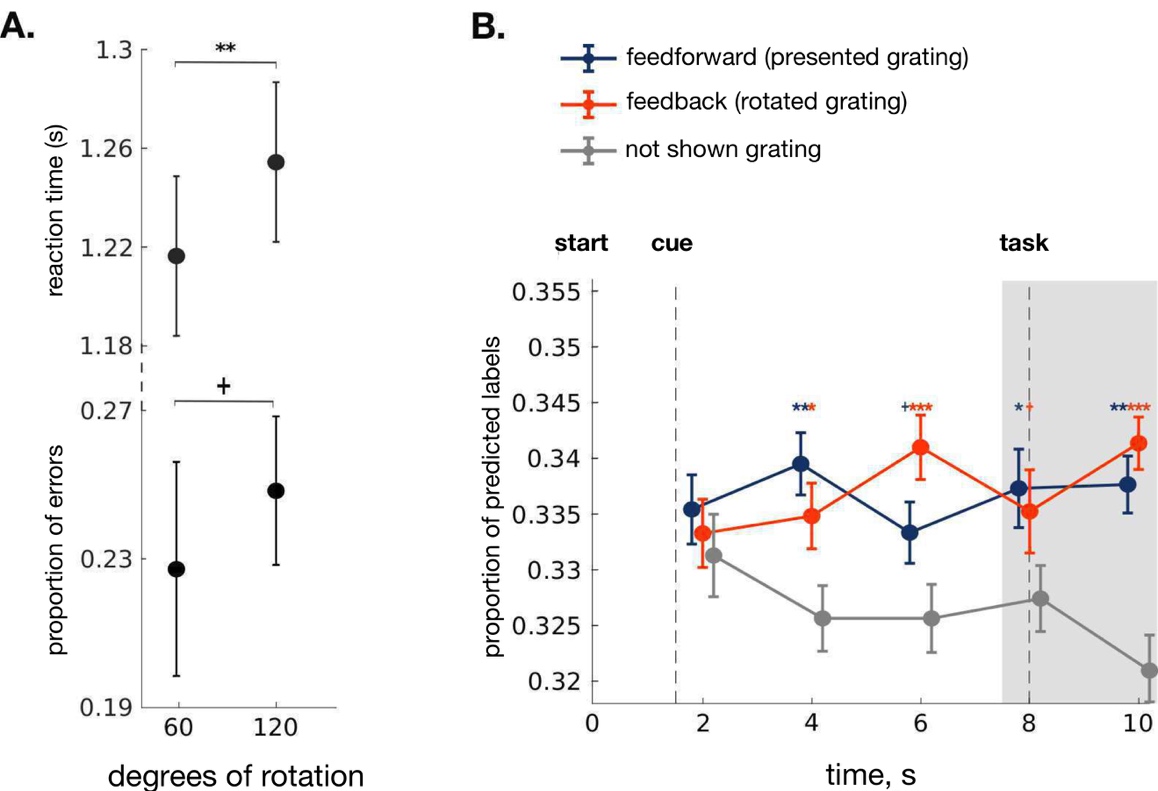

**Supplementary Figure 1.** Behavioral results and classifier decisions in V1. **A.** Reaction time was shown to be slower and error rate to be higher when responding to the probe after 120° mental rotation compared to 60° ^1^. +: p<0.07, **: p < 0.01. **B.** To establish the concurrent representation of perceived and mentally rotated contents in V1 we averaged classifier predictions across the cortical depths. This yielded three time courses, reflecting how often classifiers predicted the orientation of the presented, the rotated and the unused gratings. To estimate the presence of perceptual representations, we ran a 2×2 repeated measures ANOVA with factors Signal Type (perceived vs. unused grating) and Time Points (4,6,8, and 10 seconds after onset). In this time interval, perceptual representations were evident from a significant main effect of Signal Type (Figure 2B, F_1,66_=15.3, p=0.0007). To estimate the presence of feedback signals, we ran a similar ANOVA, now comparing the proportion of predictions for the rotated grating and the unused gratings. We also obtained a significant main effect of Signal Type for the mentally rotated contents (Figure 2B, F_1,66_=21.55, p=0.0001). We also provide the results of the t-tests for every time point (uncorrected for multiple comparisons). Both perception and rotation signals were similar across the time points examined (interaction with time, both F<1.27). When comparing representations of perceived and rotated contents, no significant differences were found (F=0.16), also when considering individual time points (all t<1). These results establish the presence of feedforward and feedback signals in V1. All error bars denote standard error of the mean over subjects. +: p<0.09, *: p<0.05, **: p < 0.01, ***: p < 0.001.

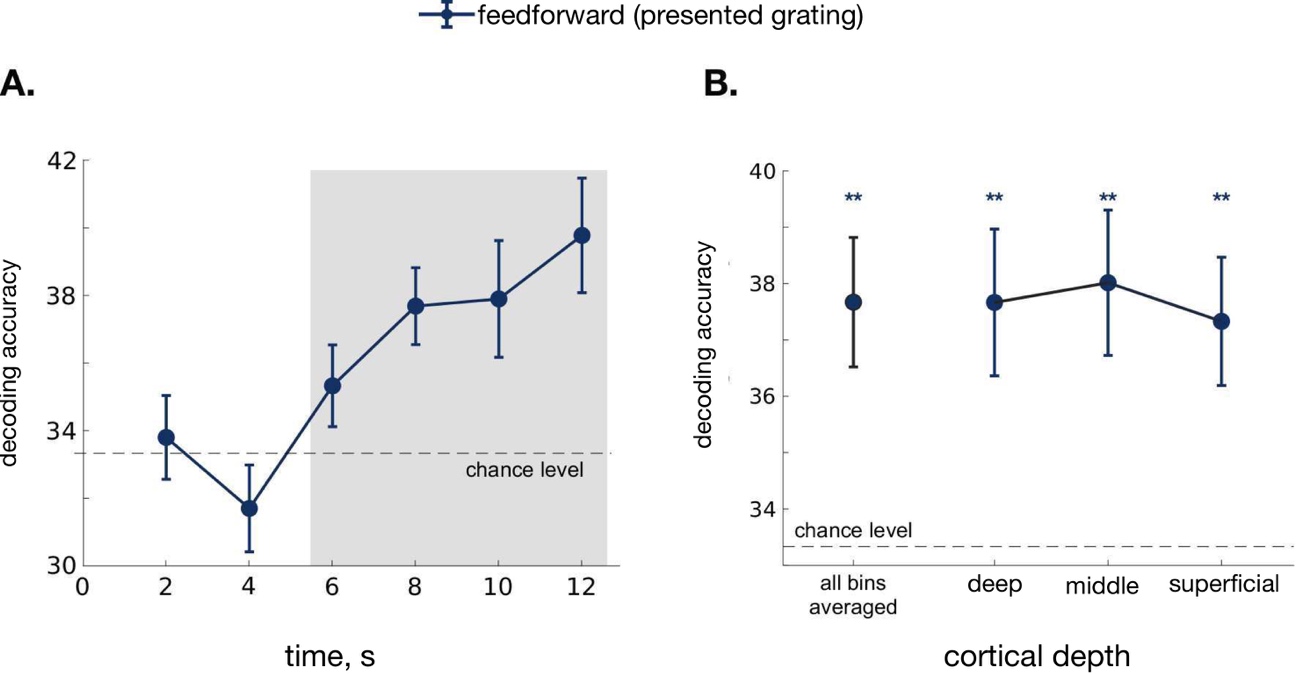

**Supplementary Figure 2.** **Decoding orientation information from the perceptual localizer task in V1.** **A.** The time series of perception signal averaged across all grey matter bins**.** In order to establish the representation of perception signal in the localizer task, we trained a three-class classifier to differentiate between the three grating orientations using all the trials in the run leaving one trial out. Start denotes the start of the 12 seconds trial in the perceptual localizer task.  The shaded area highlights the time points selected for the further depth-specific analysis. **B.** Classifier decisions over the measurements at 6, 8, 10 and 12 seconds after the trial onset across the cortical depth bins reveal robust neural information about the feedforward perceived orientation (F_1,66_ = 909.68, p=2.2e-19). We also found significant representation of orientation information in each depth bin separately (for deep: t_22_=3.33, p=0.0015, middle: t_22_=3.62, p=0.0015, superficial: t_22_=3.51, p=0.0015); however, no difference between the different bins was found.  Horizontal line represents a chance level (33,33%) against which the decoding accuracy estimates were tested. All error bars denote standard error of the mean over subjects. *: p<0.05, **: p < 0.01, ***: p < 0.001.

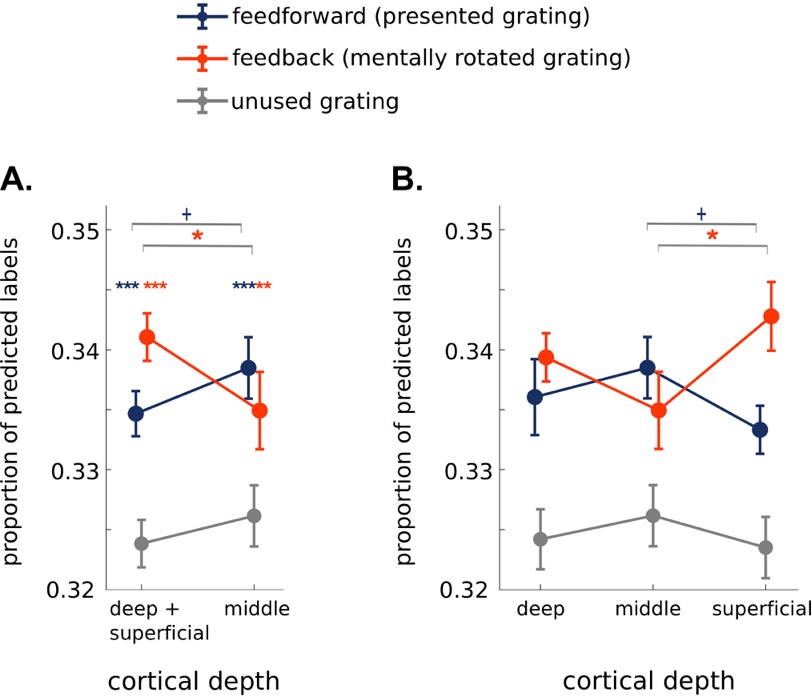

**Supplementary Figure 3. Classifier decisions in V1 at an extended time interval (6, 8 and 10 sec.) after trial onset.** Since in our experiment we find a representation of mental rotation already at 6 seconds after rotation cue, we also provide the in-depth analysis for the time interval comprising measurements at 6, 8 and 10 seconds. **A.** At the time interval spanning 6,8 and 10 seconds, we found a strong representation of mental rotation contents in the outer cortical bins (t=5.4, p=1*10^-5, Cohen’s d=0.99) and a trend in the middle bin (t=1.7, p=0.05, Cohen’s d=0.35). The feedforward signal was significantly above unused grating in all the cortical bins (at the middle depth: t=3.1, p=0.008, Cohen’s d=0.65; at the outer depths: t=3.6, p=0.0008, Cohen’s d= 0.75), in accordance with the results at the initially chosen time interval in Fig.2A.  The 2x2 Depth-by-Signal interaction was at the level of trend (F=3.4, p=0.078). Specifically, the outer cortical bins carried mental rotation representation compared to the middle depth (t=1.8, p=0.04, Cohen’s d= 0.38) and a trend towards more strongly represented perception signal than mentally rotated gratings was found at the middle cortical depth (t=1.6, p=0.06, Cohen’s d= 0.32). Additionally, a direct comparison of the two signal types in the outer cortical bins revealed more strongly represented mentally rotated contents compared to the perceived ones (t=1.8, p=0.04, Cohen’s d= 0.38) indicating an exclusive role of outer cortical depths in carrying feedback signal. **B.** Separate analysis across the three cortical depths revealed a significant 2x2 Depth-by-Signal interaction between the middle and superficial cortical bins (F=7.1, p=0.01), with superficial depth carrying stronger mental rotation representation in comparison to middle depth (t=2.3271, p=0.04, Cohen’s d=0.5) and middle depth predominantly representing feedforward signal (t=2, p=0.07, Cohen’s d=0.4). Additionally, in the superficial bin we found a stronger representation of mentally rotated contents than perceived ones (t=2.24, p=0.02, Cohen’s d=0.47) confirming the predominant role of superficial cortical bin in supporting mental rotation feedback. The 2x2 Depth-by-Signal interaction between the middle and deep cortical bins as well as the 3x2 Depth-by-Signal interaction did not reach significance. Overall, including three time points in the in-depth analysis gives qualitatively similar Signal-by-Depth dissociation as in the main analysis (Fig.2) where mentally rotated contents are more strongly represented in the outer bins, mainly at the superficial depth, and perception contents are present across all the three depths. All error bars denote standard error of the mean over subjects. +: p<0.09, *: p<0.05, **: p < 0.01, ***: p < 0.001.

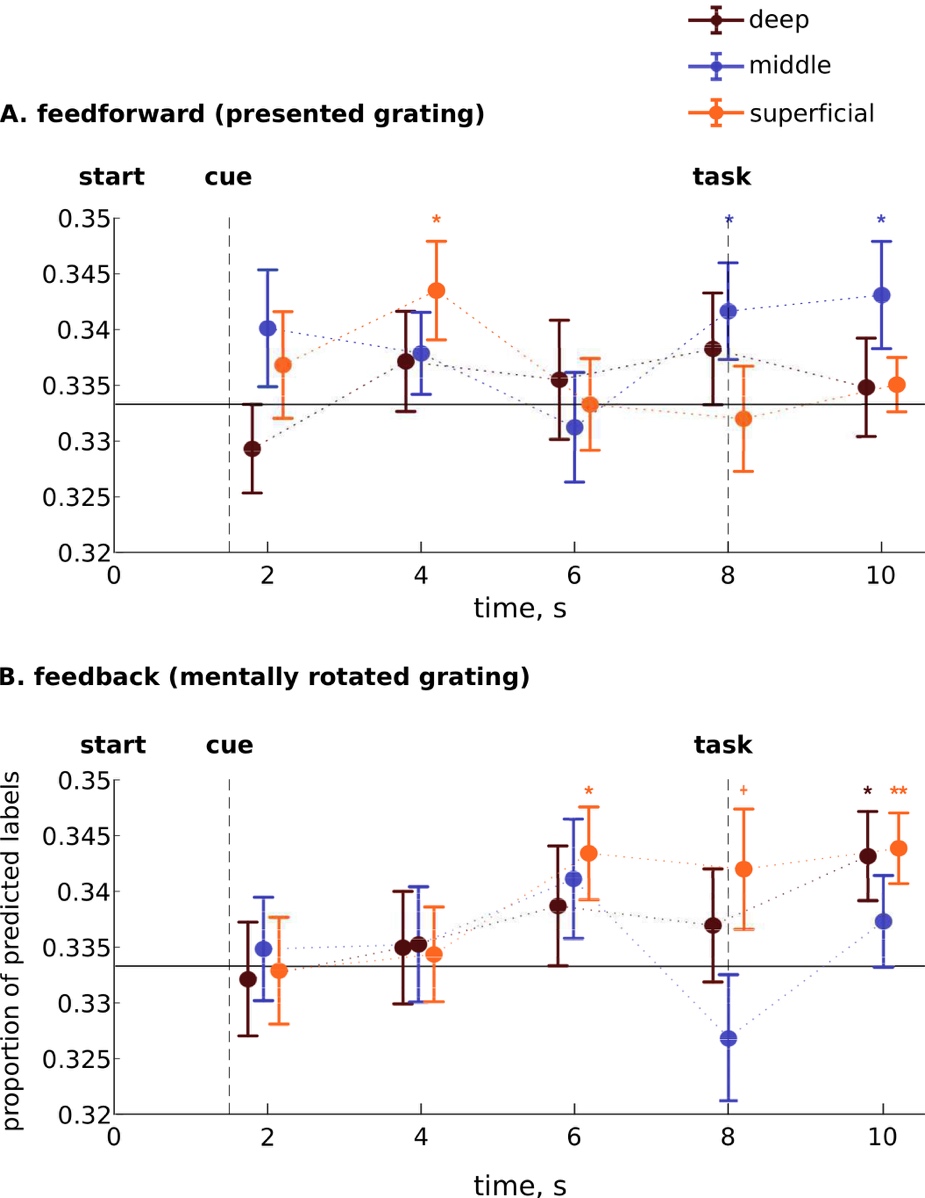

**Supplementary Figure 4.** **Depth-specific time series of classifier decisions in area V1 plotted separately for feedforward (A) and feedback signals (B).** As in the main analysis, classifier decisions for the perceived and rotated orientation were tested against decisions for the unused orientation (not shown here to avoid clutter). Grey area highlights the time interval chosen for the main analysis. All error bars denote standard error of the mean over subjects. *: p<0.05, **: p < 0.01, ***: p < 0.001 (uncorrected for multiple comparisons).

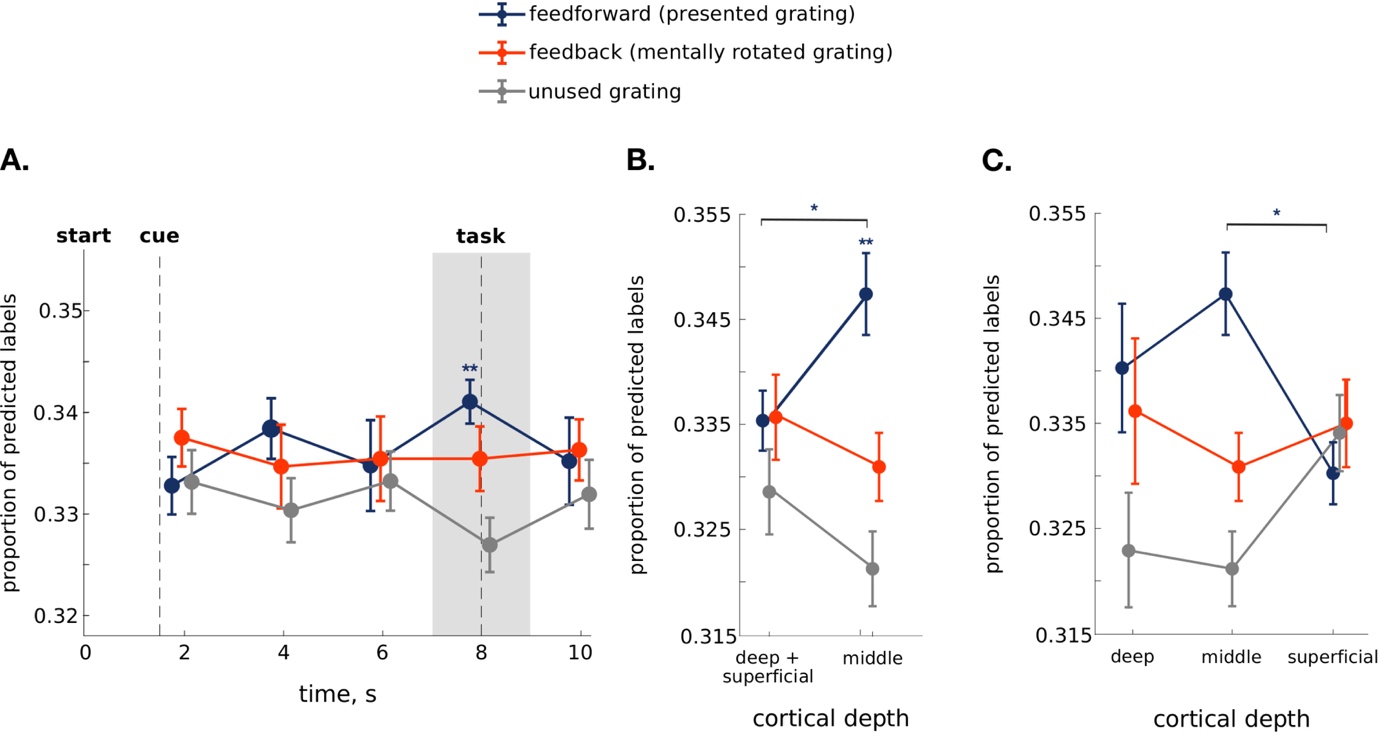

**Supplementary Figure 5.** **Classifier decisions for mental rotation and perception information in area V2.** **A.** Analysis of time courses of classifier estimates averaged across cortical depths, revealed a significant decoding of visual perception contents in area V2 measured at 8 seconds after the trial onset (t_22_=3.83, p=0.002). We found no significant decoding of mentally rotated contents in area V2 and V3. The shaded area denotes the time point selected for the further depth-specific analysis. **B.** Classifier decisions over the time interval from 6 to 8 seconds after the trial onset for the presented, mentally rotated and unused gratings in the middle and outer depths (averaged superficial and deep). The analysis revealed significant representation of the perceived contents at the middle cortical depth (t_22_=3.9, p=0.0012) compared to the unused grating. Furthermore, a stronger perception signal was found in the middle cortical bin compared to the outer bins (t_22_=2.24, p=0.035). **C.** Analysis across all the three cortical depths separately revealed a more strongly represented feedforward signal at the middle depth compared to the superficial one (p<0.05). All error bars denote standard error of the mean over subjects. *: p<0.05, **: p < 0.01.

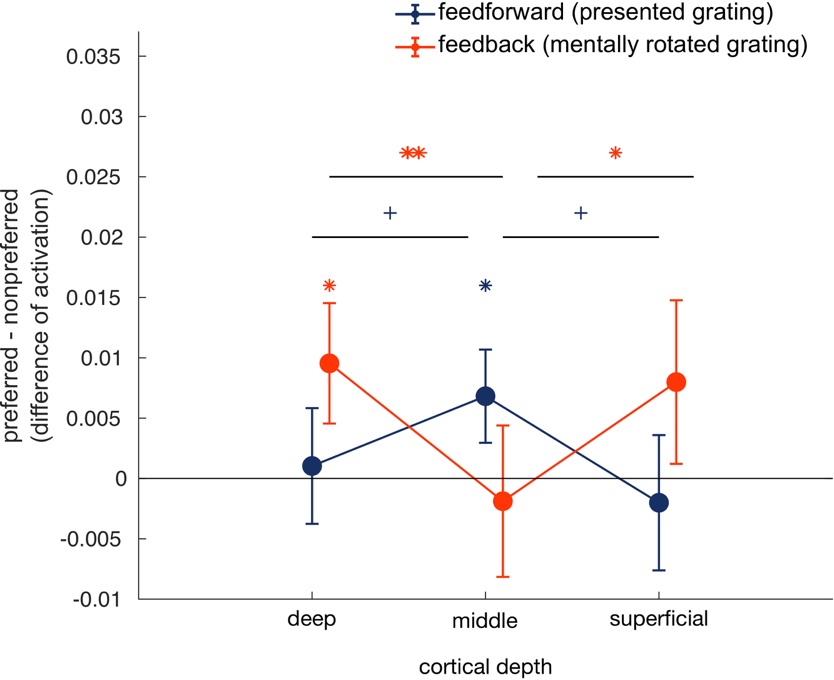

**Supplementary Figure 6. Signal-by-Depth interaction analyzed with a univariate approach.** We sorted the voxels in each cortical depth by their preference towards each of the three grating orientations based on the voxel activations in the orientation localizer task. We then picked the 300 most orientation-preferring voxels for each of the three orientations. To quantify the rotation signal, we then calculated the difference between the activation for rotated grating and two other gratings (non-preferred) within the voxels which preferred the rotated grating condition. To quantify the perception signal, we calculated the difference between the activation for perceived grating and two other gratings within the voxels which preferred the perceived grating condition. Prior to the analysis, the voxel responses both in the orientation localizer task and in the main experiment were high-pass filtered using spm12 (but not for the multivariate analysis (see Methods)). We found a signal X depth interaction (3X2 interaction: F(2,44)=3.34, p=0.04; 2X2 interaction: F(1,22)=6.5, p=0.02), showing stronger rotation information in the superficial and deep cortical bins and stronger perception information at the middle depth. In detail, perception signal was significantly above 0 at the middle cortical depth (t(22)=1.8, p=0.05, Cohen’s d=0.37) and trended to be stronger than in the outer bins (middle vs. deep bins: t(22)=1.6, p=0.06 Cohen’s d=0.33; middle vs. superficial bins: t(22)=1.6, p=0.06, Cohen’s d=0.34). The rotation signal was significantly above 0 in the deep bin (t(22)=1.9, p=0.03, Cohen’s d=0.4) and stronger in the outer bins than the middle bin (middle vs. deep bin: t(22)=2.6, p=0.007, Cohen’s d=0.55; middle vs. superficial bin: t(22)=1.8, p=0.04, Cohen’s d=0.37).
